# Monkeybread: A Python toolkit for the analysis of cellular niches in single-cell resolution spatial transcriptomics data

**DOI:** 10.1101/2023.09.14.557736

**Authors:** Matthew N. Bernstein, Dillon Scott, Cynthia C. Hession, Tim Nieuwenhuis, Jacqueline Gerritsen, Siamak Tabrizi, Vandana Nandivada, Matthew A. Huggins, Meixue Duan, Shruti Malu, Ming Tang

## Abstract

Spatial transcriptomics technologies enable the spatially resolved measurement of gene expression within a tissue specimen. With these technologies, researchers can investigate how cells organize into cellular niches which are defined as distinct regions in the tissue comprising a specific composition of cell types or phenotypes. While general-purpose software tools for the exploratory analysis of spatial transcriptomics data exist, there is a need for tools that specialize in the analysis of cellular organization into niches. This can further enhance the downstream application of these data towards drug target discovery, target validation, and biomarker development. We present Monkeybread: A Python toolkit for analyzing cellular organization and intercellular communication in single-cell resolution spatial transcriptomics data. We applied Monkeybread to a human melanoma sample to demonstrate its utility in identifying cellular niches with diverse immunogenic compositions in the tumor microenvironment. We found that these niches were differentially enriched for immunogenic and tolerogenic macrophage populations that could be correlated to T cell abundance. These findings highlight how Monkeybread can be used for revealing underlying biology of the tumor microenvironment, and in the future, for understanding the influence of these niches on response to available treatments and discovery of novel drug targets.

## Introduction

Spatial transcriptomics technologies enable the measurement of gene expression while resolving the spatial locations of those measurements within the tissue specimen. With these methods, researchers can study how gene expression correlates with cellular organization and intercellular communication in the tissue microenvironment. To unleash their potential, a variety of software tools have been developed to facilitate the analysis of the data they generate. Many of these tools address computational challenges inherent in the analysis of data generated by technologies that measure gene expression at the resolution of spatial *spots* that are locations in the tissue containing multiple cells (ranging from a few to tens of cells) from which a gene expression measurement was made (such as data generated by the 10x Visium platform). For example, tools have been developed to deconvolve the abundance of specific cell types within each spot ^1–4^ or to cluster spots in a spatially-aware manner ^5,6^.

However, fewer tools address the challenges inherent to analyzing single-cell resolution spatial transcriptomics data, such as those generated by multiplexed error-robust fluorescence in situ hybridization (MERFISH), which often generates data from hundreds of thousands of spatially resolved individual cells. First, when visualizing the full tissue specimen, there often exist so many cells that it becomes difficult to visually discern distributions of cell types (especially rare cells) or gene expression patterns. Furthermore, while methods exist for identifying broad tissue regions using spatially aware clustering of gene expression profiles ^5–7^, except for SPIAT ^8^ (which is written in R), we are not aware of any tools for the Python ecosystem that identify cellular niches as defined by cells that cluster according to the composition of the user-defined cell types in their spatial neighborhoods. Understanding the spatial cellular niches that comprise the tissue microenvironment is essential for understanding basic physiology ^9–11^ and disease ^12–14^. Lastly, though some methods, such as squidpy ^15^, stLearn ^16^, Seurat ^17^, Giotto ^18^, and Voyager ^7^ provide broad functionality for analyzing spatial transcriptomics data, these methods are generalized to work on spatial data at both the spot-level and single-cell level but don’t address the aforementioned challenges by specializing in the analysis of cellular niches in single-cell resolution spatial data.

In this work, we present Monkeybread: A Python toolkit that provides broad functionality for analyzing and visualizing cellular organization, cellular niches, and intercellular communication via ligand-receptor interactions. Monkeybread is designed to operate on spatial datasets stored in memory as AnnData objects, and thus, can be integrated into workflows that use scanpy ^19^, squidpy, or other tools in the scverse ecosystem ^20^. To provide a case-study into how Monkeybread can be used, we applied it to a human melanoma sample from Vizgen’s immuno-oncology public data release ^21^. This analysis revealed cellular niches with diverse immunogenic repertoires in the tumor microenvironment (TME).

## Results

### Monkeybread provides a suite of tools for exploring cellular organization and communication in single-cell spatial transcriptomics data

Monkeybread enables investigations into how cells organize into cellular niches, here defined as regions within the tissue microenvironment comprising a specific composition of cell types or phenotypes. To do so, Monkeybread first selects the neighboring cells of each cell (those that fall within a pre-specified radius of each cell) and counts the number of cells of each cell type or phenotype to form an *N × C* neighborhoods matrix, ***Y***, where ***Y*** is the number of cells and *N* is the number of cell type or phenotype labels. Element *i, j* represents the number of neighbors of cell *i* that are of cell type *j*. To account for neighborhoods with differing numbers of cells, Monkeybread provides the option of normalizing these neighborhood counts by dividing each cell’s neighbor counts by its total number of neighbors. To account for the large differences in cell abundances between cell types, Monkeybread also provides the option of computing the z-scores of each neighborhood count (or normalized count), thus placing count values between cell types on similar scales. Finally, niches are computed by clustering the rows (corresponding to cells) of the normalized neighborhoods matrix. By default, Monkeybread uses the Leiden clustering algorithm ^22^.

Monkeybread provides a lightweight statistical test for testing enrichment of cellular colocalization between pairs of cell types (Figure S1). This test considers the following test statistic: for two cell types *A* and *B* under test, for each cell of type *A*, we find the distance to its nearest cell of type *B*. We then count the number of such shortest distances from cells of type *A* to type *B* that fall below some user-provided threshold distance *D*. To test for significance, a permutation test is performed that generates a null distribution of the test statistic by permuting the spatial coordinates of non-type *A* cells. This test can optionally be run on cells within specific niches to condition the analysis on co-localization patterns explained by the organization of the tissue into cellular niches.

When visualizing the full tissue specimen, there often exist so many cells that it becomes difficult to visually discern the spatial distribution of cells of a given cell type, especially rare cell types, or those expressing a given gene. Monkeybread provides functionality for visualizing and comparing the spatial density of cells with a specific cell type label or phenotype across the tissue. More specifically, Monkeybread uses kernel density estimation to visualize regions with a high concentration of cells exhibiting rare or specific phenotypes (Methods). In addition, Monkeybread provides functions built atop scanpy for easily zooming in on specific regions of the tissue and sub-setting the data to the zoomed-in regions within the smaller field of view.

Lastly, Monkeybread implements a lightweight statistical test developed by He *et al*. (2021) ^23^ for identifying intercellular communication via ligand-receptor binding (Methods; Figure S2). This method is related to a spatial ligand-receptor co-expression test implemented in Giotto but here is implemented in Python. It differs from that implemented by Squidpy in that Squidpy’s uses CellPhoneDB ^24^, which does not consider spatial proximity between cell types. In addition to this test, Monkeybread provides a visualization tool, built atop matplotlib ^25^ and scanpy ^19^, that depicts which pairs of cells in the tissue may be communicating via a given ligand-receptor by drawing a line between each pair of neighboring cells of types *A* and *B* that are expressing the given ligand and receptor. These lines are colored according to the magnitude of their ligand-receptor co-expression scores. This visualization tool enables the user to quickly see where in the tissue neighboring cells are co-expressing the ligand-receptor pair.

### Monkeybread identifies spatial regions enriched for immunogenic and tolerogenic macrophages in a human melanoma tumor

We applied Monkeybread to a human melanoma sample from Vizgen’s immuno-oncology public data release ^21^ to study the cellular organization of the TME. First, we annotated the cells with both a coarse and granular set of cell type labels by first clustering the cells and manually cross-referencing the genes expressed in each cluster with known marker genes (Figure 1a,b, S3-7; Methods). We examined the spatial distribution of the broad cell types using Monkeybread’s tool for visualizing cell spatial densities and found malignant cells occupying the core of the tumor surrounded by regions of dense stromal and immune cells (Figure 1.c).

**Figure 1.**
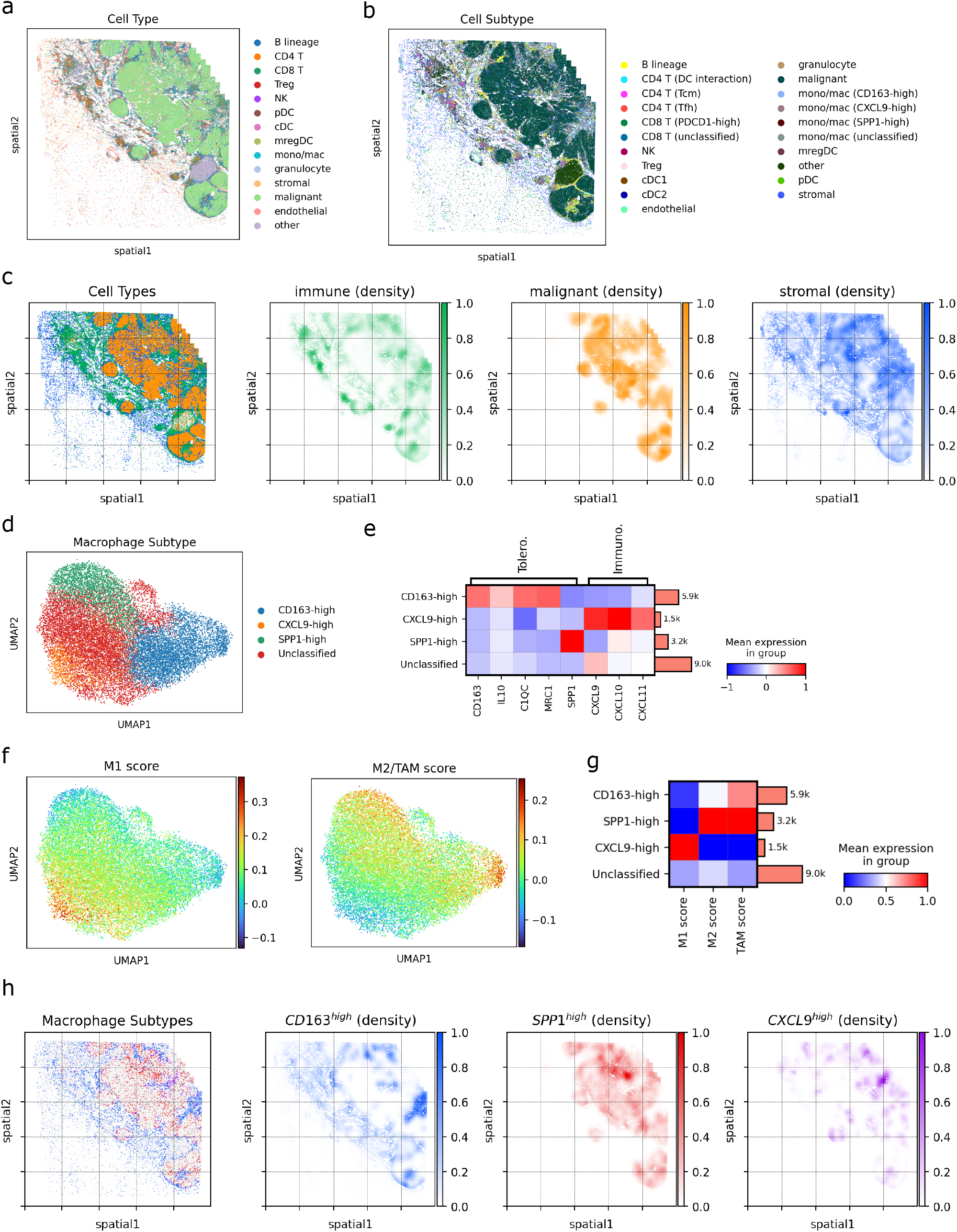
Spatial distribution of macrophage populations in a human melanoma. **(a)** Annotated broad cell types and **(b)** granular cell types of cells in the human melanoma tumor dataset. **(c)** Locations and densities of immune, malignant, and stromal cells. The density plots were generated with Monkeybread’s density visualization tool. **(d)** UMAP plot of all annotated macrophages colored by their annotated subtype. **(e)** Mean normalized expression (with z-score transformation) of known tolerogenic and immunogenic marker genes within each macrophage subtype. **(f)** UMAP plot of macrophages colored by the aggregate expression score of genes found to be differentially expressed between in vitro differentiated M1-like macrophages versus in vitro differentiated M2 and TAM-like macrophages. Scores were computed with scanpy’s “score_genes” function. **(g)** Heatmap depicting the aggregate M1, M2, and TAM expression score in each macrophage subpopulation. Columns were standardized such that the maximum value was set equal to 1 and minimum value was set equal to zero. **(h)** Locations and densities of the annotated macrophage subpopulations. The density plots were generated with Monkeybread’s density visualization tool.

Within the monocyte/macrophage compartment, we identified three distinct populations (Figure 1d). The first displayed a putatively immunogenic phenotype as indicated by high expression of CXCL9, CXCL10, and CXCL11 ^26^. The second displayed markers commonly associated with a tolerogenic, M2-like phenotype (CD163 and MRC1) as well as other genes associated with immune suppression (C1QC and IL10) ^27,28^. A final population expressed SPP1, which has also been associated with immune suppression and poor clinical outcome (Figure 1.e) ^29^.

To further understand the immunological role of these macrophage populations, we compared their transcriptional profiles to macrophages differentiated from PBMC-derived monocytes *in vitro* under three conditions: M1-like macrophages were developed *in vitro* through polarization of human monocytes by IFNγ and TLR4 stimulation. This population is considered to have anti-tumor functions including release of proinflammatory cytokines, phagocytosis, and ability to cross-present extracellular antigens produced by dying tumor cells to activate CD8 T cells ^30^. Second, M2-like macrophages were developed *in vitro* through intensive IL4 stimulation of monocytes. M2 macrophages are known to promote many pro-tumorigenic processes such as angiogenesis, immune suppression, hypoxia induction, tumor cell proliferation, and metastasis ^31^. Lastly, we deployed a third population that represents a better *in vitro* model for tumor associated macrophages (TAM), developed under conditions more closely resembling the TME ^32^. We performed bulk RNA-seq on these three populations and performed differential expression (DE) analysis between the M1-derived population and the M2-derived population as well as between the M1-derived and TAM-derived populations (Methods; Table S1). We found 134 genes higher in M1 vs. M2 and 144 genes higher in M1 vs. TAM (adj. p < 0.05). We took the intersection of these two gene sets to derive an M1-signature consisting of 101 genes that we used to score the *in vivo* macrophages (Methods). Similarly, we found 80 genes higher in the M2 vs. M1 populations and 78 genes higher in the TAM vs. M1 populations (adj. p < 0.05). We took the union of these two genes sets to derive an M2/TAM-signature comprising 100 genes that we used to score the *in vivo* macrophages. Overall, we observed high expression of the M1-signature in the CXCL9-high macrophages and high M2 and TAM signatures in the CD163-high and SPP1-high macrophages (Figure 1.f,g). In summary, these results support the immunogenic function of the CXCL9-high and tolerogenic function of the CD163-high and SPP1-high macrophages in the tumor microenvironment.

Next, we examined the spatial distribution of these three macrophage populations using Monkeybread’s kernel density estimation tool and found that these cells occupy distinct regions within the tissue (Figure 1.h). Interestingly, the CXCL9-high and CD163-high populations were concentrated in regions with higher immune cell abundance (immune “hot” regions) and in regions surrounding the tumor. In contrast, the SPP1-high macrophages were found in regions with fewer immune cells (immune “cold” regions) and mostly found within the tumor in regions of dense malignant cells. Although the gene expression signatures of the CD163-high and SPP1-high macrophages indicate a suppressive function, their distinct spatial distribution indicates that they are playing further differentiated roles in the TME.

We then used Monkeybread to identify distinct cellular niches of immune cells in the tumor microenvironment (Figure 2.a). These niches included both immunologically cold (Niches 1 and 10) and hot niches (all remaining niches; Figure 2.b). Niche 7 was enriched for CXCL9-high macrophages, Niches 0 and 10 for CD163-high macrophages, and Niche 1 for SPP1-high macrophages. Niche 7 contained a higher enrichment of diverse immune cells than Niches 0, 1, and 10. We then examined the mean expression in immune cells within each niche of a ten-gene interferon-gamma related signature that has been shown to correlate with response to immunotherapy ^33^ (Fig. 2.c). Niche 7 displayed high overall expression of genes in this signature while Niche 10 displayed markedly low expression. Moreover, Niches 0 and 10 (the CD163-high enriched niches) displayed marked lower levels of IFNG than the other niches. Together, these results indicate that Niche 7, and in turn CXCL9-high macrophages, have an immunogenic role whereas CD163-high macrophages may have a suppressive role in the TME.

**Figure 2.**
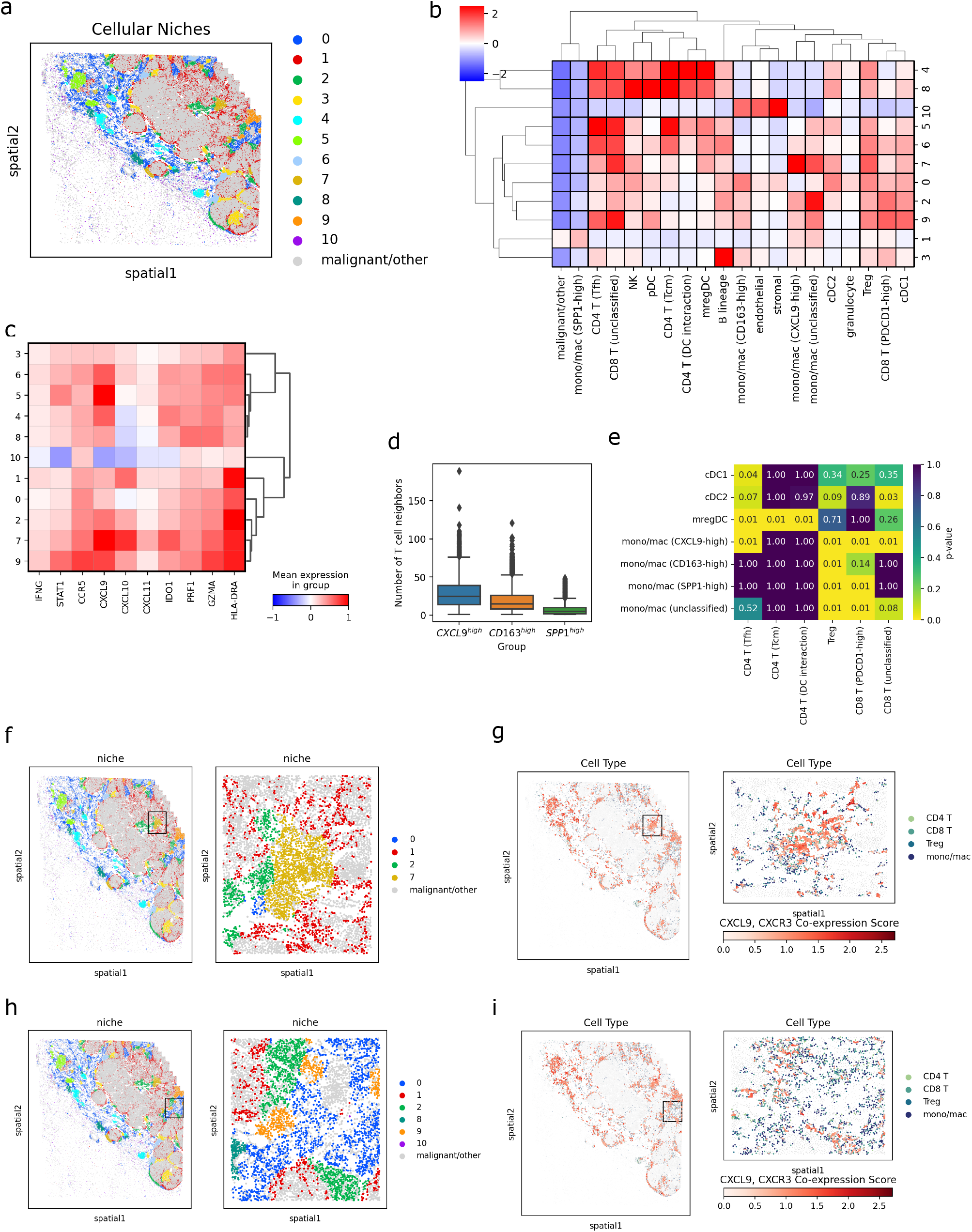
Analysis of cellular niches and intercellular communication in a human melanoma. **(a)** Immune cellular niches in a human melanoma tumor dataset identified by Monkeybread. **(b)** Heatmap displaying the mean neighborhood enrichment scores for each cell type within each cellular niche. **(c)** Mean normalized expression (with z-score transformation) within each niche of a 10-gene interferon-response related signature from ^33^. **(d)** Distribution over the number of neighboring T cells (within a radius of 75 μm) surrounding each macrophage subtype. **(e)** Heatmap displaying the p-values from Monkeybread’s cell type colocalization test between all myeloid cell subtypes and T cell subtypes. **(f)** Plot generated by Monkeybread zooming in to a region annotated as Niche 7. **(g)** In this same region, Monkeybread was used to plot macrophage-T cell pairs co-expressing CXCL9 and CXCR3. **(h**,**i)** Similar plots to those shown in (f) and (g), but zooming in to a region annotated as Niche 0.

Next, we used Monkeybread to plot the distribution of the number of T cells that are neighbors of CXCL9-high, CD163-high, and SPP1-high macrophages. Consistent with the niche analysis, we found that the CXCL9-high macrophages are surrounded by the most T cells whereas SPP1-high macrophages are surrounded by the fewest (Figure 2.d). Next, we ran Monkeybread’s cell colocalization statistical test between all pairs of myeloid cell subtypes and all T cell subtypes. We found that CD163-high and SPP1-high macrophages only displayed enriched co-localization with T regulatory (Treg) cells and PDCD1-high CD8 T cells whereas CXCL9-high macrophages also colocalized with T follicular helper cells (Tfh) cells and unclassified CD8 T cells (Figure 2.e). These results were consistent with the niche analysis, which showed higher Tfh enrichment in Niche 7 than in Niches 0, 1, or 10. This is a highly relevant observation since Tfh cells have been shown to correlate with reduced tumor growth and improved survival in multiple solid tumors types ^34^. Together these results further support that these macrophage populations play differing immunological roles within distinct niches in the TME.

Lastly, we used Monkeybread to test for co-expression of the ligand receptor pairs CXCL9/CXCR3 and CXCL10/CXCR3 between all macrophages and their neighboring conventional T cells and found significant co-expression of CXCL9 and CXCR3 (p = 0.001) as well as between CXCL10 and CXCR3 (p=0.040). Using Monkeybread’s function to visualize this co-expression, we confirmed higher co-expression of CXCL9/CXCR3 in Niche 7, which is enriched for CXCL9-high macrophages (Figure 2.f,g), compared to Niche 0, which is enriched for CD163-high macrophages (Figure 2.h,i).

## Discussion

We present Monkeybread: A Python toolkit to facilitate the analysis of single-cell resolution spatial transcriptomics data enabling investigators to explore how cells organize into cellular niches in the tissue microenvironment. Monkeybread operates on data represented as AnnData objects and thus, can be integrated into analyses that utilize scanpy, squidpy, and other tools in the scverse ecosystem. We used Monkeybread to explore the cellular organization in a spatial transcriptomics dataset from a human melanoma tumor. This analysis demonstrated how Monkeybread can be integrated with scanpy, and other Python analysis packages, to explore the tissue microenvironment. Specifically, we identified three different macrophage populations with putatively different functional roles and, with Monkeybread, found these populations participate in distinct immunological cellular niches in the TME. Altogether, we expect Monkeybread to be a useful complement to existing Python packages designed for the exploration of single-cell spatial transcriptomics data.

### Limitations of the study

There are several avenues that would benefit from further investigation. First, Monkeybread’s computational workflow for identifying cellular niches requires the pre-specification of a clustering parameter that determines the number of clusters of cell type neighborhood profiles. Though methods have been proposed for selecting the number of clusters in single-cell transcriptomics data ^35–37^, to our knowledge, no work has addressed the specific problem of determining the number of clusters to use when clustering cell type neighborhood profiles when identifying cellular niches. Nonetheless, general visualization tools for assessing cluster granularity, such as clustering trees ^38^, will be explored as additions to Monkeybread’s visualization toolkit.

Second, Monkeybread only facilitates exploratory analysis of the gene expression matrix that is downstream of image processing, cell segmentation, and cell type annotation that are prerequisite steps in the analysis of imaging-based spatial assays. Errors in these steps may propagate into the analyses that Monkeybread can perform.

Third and finally, future work will entail extending the case-study, which focused on the analysis of a single melanoma tumor sample, to more datasets and tumor types. In this case study, we identified diverse cellular niches of immune cells in the tumor microenvironment and these niches were differentially enriched for immunogenic and tolerogenic macrophages. Future work will expand this analysis to more datasets with a focus on understanding the similarities and differences between immunological niches between tumors.

## Author contributions

M.N.B and M.T. conceived the project and wrote the first version of the manuscript. D.S. and M.N.B. implemented the software. M.T. and M.D. tested the software. S.T., V.N., J.G., and C.C.H. executed the macrophage differentiation experiment and prepared the sequencing library. T.N. analyzed the bulk RNA-seq data. M.N.B. performed the analysis of the human melanoma dataset. M.A.H. and S.M. provided domain expertise for the analysis of the human melanoma data.

## Declaration of interests

All authors were paid employees of Immunitas Therapeutics, Inc. at the time this work was performed.

## Methods

### Visualizing cell densities

Monkeybread uses kernel density estimation to visualize the spatial distribution of cells that exhibit a certain phenotype of interest (*e*.*g*., T cells), which smooths out the discrete cell locations enabling easier visualization of regions with high or low density ^39^. Specifically, let ***s*** ∈ ℝ^2^ denote the spatial coordinates of a cell exhibiting the target phenotype. We assume that there exists an underlying probability distribution *p*(***s***) over ***s*** that we seek to estimate via kernel density estimation. To visualize the concentration of cells across the tissue, we seek to color each cell by the value of this probability density function at its given coordinate. Let ***s***_1_ … ***s***_*C*_ ∈ ℝ^2^ be the observed spatial coordinates in the data of the *C* cells that exhibit the target phenotype. Then, *p*(***s***) is estimated via,

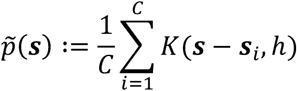

where *K* is a Gaussian kernel with bandwidth parameter *h*,

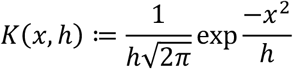

We then transform the kernel density estimates to fall between zero and one. Specifically, for cell *j*, its color intensity would be computed via

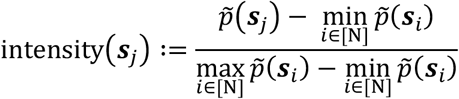

where *N* is the total number of cells in the dataset and [*N*] are the set of integers from 1 to *N*.

To speed up computation, Monkeybread provides the option to run grid-based approximation ^40^. Specifically, a grid is overlayed onto the spatial data and each cell is assigned to the grid space that it falls within. Let *G*_1_, …, *G*_*M*_ be the partitioning of cells into *M* grid spaces and let 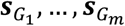 ∈ ℝ^2^ be the coordinates of their centroids. Because cells within the same grid share the same “collapsed” coordinates, the kernel density estimate for a given location requires iterating over all grid spaces rather than all cells:

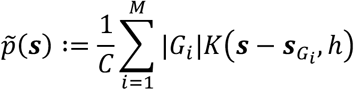

In the limit, as the grid becomes fine-grained enough such that each grid space contains a maximum of one cell, this approximation will converge on the true kernel density estimate. A further speedup is made by coloring each cell according to the kernel density estimate at its grid space’s centroid rather than its own coordinate – that is, the density estimate is calculated at each grid’s centroid and mapped back to each cell within that grid space. Lastly, to derive the color intensity used in the visualization, we transform these density values so that they fall between zero and one. Specifically, let *g*(*j*) be the function that maps each cell *j* to the grid space that it falls within. Then, for cell *j*, its color intensity is computed as

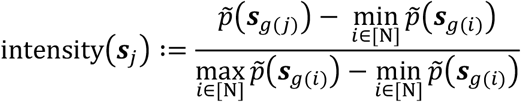

where *N* is the total number of cells in the dataset.

### Spatial ligand-receptor co-expression test

Monkeybread implements the statistical test for spatial co-expression described by He *et al*. (2021) ^23^. To review, this test considers the following test statistic: for a given ligand, *L*, and receptor, *R*, and two cell types, *A* and *B*, under test, for each cell of type *A*, we first identify the neighboring cells of type *B*. We then compute the following ligand-receptor co-expression score:

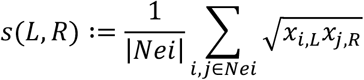

where *Nei* is the set of all pairs of neighboring cells between types *A* and *B, x*_*i,L*_ is the expression of gene *L* in cell *i* (of cell type *A*), and *x*_*j,R*_ is the expression of gene *R* in cell *j* (of cell type *B*). Statistical significance of this score is assessed via a permutation test in which the pairings of neighboring cells are repeatedly permuted, and the test statistic is recalculated to form the null distribution.

### Preprocessing and cell type annotation

The melanoma tumor dataset was downloaded from Vizgen’s website at https://vizgen.com/data-release-program and preprocessed with scanpy. Specifically, cells with less than 10 transcript counts were filtered out of the dataset resulting in a total of 189,071 cells. Transcript counts were normalized via log (*c*/*s* + 1) where *c* is the transcript count for a given gene in a given cell and *s* are the total transcript counts in that cell. Before clustering, PCA was performed with 50 components and the k-nearest-neighbors (KNN) graph was computed on the principal components using k=15 neighbors. Cell types were annotated by first clustering the full set of cells using the Leiden clustering algorithm ^22^ on the KNN graph and then labeled according to their expression of known marker genes within each cluster (Figure S4). Clusters that expressed multiple marker genes (indicating potential heterogeneity with respect to their constituent cell types) were further sub-clustered and annotated (Figures S5-7).

### In vitro differentiation of M1, M2, and TAM-like macrophages

Healthy Donors PBMNCs were acquired and CD16 negative monocytes (M0) were isolated with the Miltenyi biotec kit, (#130-117-337) following the manufacturer’s instructions. To develop polarized macrophages, monocytes were seeded on 100mm culture dishes with X-Vivo15 media supplied with 5% human serum. To generate M1 macrophages, cells were cultured for 5 days under GM-CSF stimulation and then for 2 days with IFNg and given a final 18h touch of LPS treatment. M2-like macrophages were developed through 5 days of culture with M-CSF and further 48h treatment of IL-4 stimulation. TAM development was accomplished via a continuous 7-day of cell culture under the influence of M-CSF+IL4 and IL10. The medium was enriched with 20% tissue conditioned medium (TCM) medium as the required supplement. Through the last 48h of culture, TGFb stimulation was added to the culture medium as well. Medium change for the culture systems was administrated once during the 7 days of culture. Tissue conditioned medium (TCM) was generated from MDA MB-231 breast cancer cell line following the protocol described by Benner et al. (2019) ^32^.

### Bulk RNA-seq

Cells derived from two patient donors were cultured in preparation for bulk RNA sequencing as described above, with four *in vitro* conditions per donor (monocyte control, M1 macrophages, M2 macropages, and TAM). After harvesting and counting, 1 million cells were aliquoted per culture condition and centrifuged at 400g for 5 minutes. Pelleted cells were resuspended in 350 uL Buffer RLT (Qiagen) + 1% BME, and vortexed for 30 seconds to 1 minute to disrupt the cells. Lysates were then frozen at -80°C and stored until ready to proceed with RNA isolation.

Total RNA was isolated from cell lysates using the RNeasy Mini Kit (Qiagen) per the “Purification of Total RNA from Animal Cells using Spin Technology” section of the kit manual. An optional DNase I treatment was included and performed using the RNase-Free DNase Set (Qiagen). Total RNA concentration was evaluated by RNA HS Qubit (ThermoFisher), followed by RNA ScreenTape (Agilent) to assess RNA quality. All RNA samples were of sufficient concentration and RNA Integrity (RIN) value to proceed with library construction.

Bulk libraries were prepared using the NEBNext Ultra ii Directional RNA Library Prep for Illumina Kit (New England BioLabs) per Section 1 of the manufacturer’s protocol. 500ng of total RNA was used as input for each sample, and an rRNA depletion was performed using the NEBNext Poly(A) mRNA Magnetic Isolation Module (New England BioLabs). SPRIselect beads (Beckman Coulter) were used in place of NEB Sample Purification Beads for all purification steps. For the final enrichment PCR, bulk libraries were barcoded using the NEBNext Multiplex Oligos for Illumina – 96 Index Primers Kit (New England BioLabs) and amplified for 10 PCR cycles.

Bulk library quality was assessed by DNA HS Qubit assay (ThermoFisher) to measure library concentration, and by D5000 ScreenTape assay (Agilent) to evaluate library quality and size. All libraries were of sufficient concentration and quality to then prepare for sequencing. Library molar concentrations were more precisely determined by quantitative PCR (qPCR) using the KAPA Library Quantification Complete Kit – Universal (Roche). Based on calculated molar concentrations, bulk libraries were individually diluted to 20 nM and pooled equally by volume for sequencing.

Bulk libraries were submitted to Broad Institute Clinical Research Sequencing Platform for paired-end sequencing on an Illumina NextSeq500 system. The sequencing parameters consisted of 25 basepairs for Read 1, 50 basepairs for Read 2, and 8 basepairs for Index 1, with a target sequencing depth per sample of at least 20 million reads. Following sequencing, all raw data files were transferred to Immunitas servers for data analysis.

### Bulk RNA-seq Differential expression analysis

The preprocessing steps were carried out using a Snakemake pipeline ^41,42^. Briefly, the raw FASTQ files were aligned using STAR (v2.7.10b) ^43^ to genome GRCh38.primary_assembly. genome.fa using annotation file gencode.v40.annotation.gtf and the count matrix was generated by FeatureCounts (v2.0.1) ^44^. The raw counts were imported into DESeq2 ^45^ (v1.36.0) in R (v4.2.1) and differential expression analysis was carried out following the standard procedure. Two differential expression analyses were performed to compute genes that were differentially expressed between M1-derived and M2-derived macrophages as well as between M1-derived and TAM-derived macrophages.

### Identification of cellular niches

To generate cellular niches in the human melanoma tumor, we considered all immune cells and computed their normalized cell type neighborhood profiles using Monkeybread’s “neighborhoods_profile” function. All non-immune cells were labeled as “malignant/other”. We then computed the KNN graph using 100 nearest neighbors followed by clustering, via Leiden, with a resolution of 0.25. All clusters containing less than 300 cells were labeled as “malignant/other” whereas all remaining clusters were used as cellular niches for downstream analysis. This full pipeline is implemented in Monkeybread’s “cellular_niches” function.

## Data and code availability

Monkeybread is available as an open-source Python package on GitHub at https://github.com/immunitastx/monkeybread. Code used to perform the analysis of the human melanoma dataset is available at https://github.com/immunitastx/monkeybread-case-study. Gene expression data from the in vitro differentiated macrophages are available from the Gene Expression Omnibus (GEO: GSE243175).

## Notes

https://github.com/immunitastx/monkeybread

## References

1. Elosua-Bayes, M., Nieto, P., Mereu, E., Gut, I., and Heyn, H. (2021). SPOTlight: seeded NMF regression to deconvolute spatial transcriptomics spots with single-cell transcriptomes. Nucleic Acids Res. 49, e50–e50. 10.1093/nar/gkab043.

2. Cable, D.M., Murray, E., Zou, L.S., Goeva, A., Macosko, E.Z., Chen, F., and Irizarry, R.A. (2022). Robust decomposition of cell type mixtures in spatial transcriptomics. Nat. Biotechnol. 40, 517–526. 10.1038/s41587-021-00830-w.

3. Kleshchevnikov, V., Shmatko, A., Dann, E., Aivazidis, A., King, H.W., Li, T., Elmentaite, R., Lomakin, A., Kedlian, V., Gayoso, A., et al. (2022). Cell2location maps fine-grained cell types in spatial transcriptomics. Nat. Biotechnol. 40, 661–671. 10.1038/s41587-021-01139-4.

4. Andersson, A., Bergenstråhle, J., Asp, M., Bergenstråhle, L., Jurek, A., Navarro, J.F., and Lundeberg, J. (2020). Single-cell and spatial transcriptomics enables probabilistic inference of cell type topography. Commun. Biol. 3, 565. 10.1038/s42003-020-01247-y.

5. Zhao, E., Stone, M.R., Ren, X., Guenthoer, J., Smythe, K.S., Pulliam, T., Williams, S.R., Uytingco, C.R., Taylor, S.E.B., Nghiem, P., et al. (2021). Spatial transcriptomics at subspot resolution with BayesSpace. Nat. Biotechnol. 39, 1375–1384. 10.1038/s41587-021-00935-2.

6. Hu, J., Li, X., Coleman, K., Schroeder, A., Ma, N., Irwin, D.J., Lee, E.B., Shinohara, R.T., and Li, M. (2021). SpaGCN: Integrating gene expression, spatial location and histology to identify spatial domains and spatially variable genes by graph convolutional network. Nat. Methods 18, 1342–1351. 10.1038/s41592-021-01255-8.

7. Moses, L., Einarsson, P.H., Jackson, K., Luebbert, L., Booeshaghi, A.S., Antonsson, S., Bray, N., Melsted, P., and Pachter, L. (2023). Voyager: exploratory single-cell genomics data analysis with geospatial statistics. 10.1101/2023.07.20.549945.

8. Feng, Y., Yang, T., Zhu, J., Li, M., Doyle, M., Ozcoban, V., Bass, G.T., Pizzolla, A., Cain, L., Weng, S., et al. (2023). Spatial analysis with SPIAT and spaSim to characterize and simulate tissue microenvironments. Nat. Commun. 14, 2697. 10.1038/s41467-023-37822-0.

9. Rajachandran, S., Zhang, X., Cao, Q., Caldeira-Brant, A.L., Zhang, X., Song, Y., Evans, M., Bukulmez, O., Grow, E.J., Nagano, M., et al. (2023). Dissecting the spermatogonial stem cell niche using spatial transcriptomics. Cell Rep. 42, 112737. 10.1016/j.celrep.2023.112737.

10. Lu, Y., Liu, M., Yang, J., Weissman, S.M., Pan, X., Katz, S.G., and Wang, S. (2021). Spatial transcriptome profiling by MERFISH reveals fetal liver hematopoietic stem cell niche architecture. Cell Discov. 7, 47. 10.1038/s41421-021-00266-1.

11. Baccin, C., Al-Sabah, J., Velten, L., Helbling, P.M., Grünschläger, F., Hernández-Malmierca, P., Nombela-Arrieta, C., Steinmetz, L.M., Trumpp, A., and Haas, S. (2020). Combined single-cell and spatial transcriptomics reveal the molecular, cellular and spatial bone marrow niche organization. Nat. Cell Biol. 22, 38–48. 10.1038/s41556-019-0439-6.

12. Magen, A., Hamon, P., Fiaschi, N., Soong, B.Y., Park, M.D., Mattiuz, R., Humblin, E., Troncoso, L., D’souza, D., Dawson, T., et al. (2023). Intratumoral dendritic cell–CD4+ T helper cell niches enable CD8+ T cell differentiation following PD-1 blockade in hepatocellular carcinoma. Nat. Med. 29, 1389–1399. 10.1038/s41591-023-02345-0.

13. Zheng, Y., Carrillo-Perez, F., Pizurica, M., Heiland, D.H., and Gevaert, O. (2023). Spatial cellular architecture predicts prognosis in glioblastoma. Nat. Commun. 14, 4122. 10.1038/s41467-023-39933-0.

14. Barrozo, E.R., Seferovic, M.D., Castro, E.C.C., Major, A.M., Moorshead, D.N., Jochum, M.D., Rojas, R.F., Shope, C.D., and Aagaard, K.M. (2023). SARS-CoV-2 niches in human placenta revealed by spatial transcriptomics. Med. 10.1016/j.medj.2023.06.003.

15. Palla, G., Spitzer, H., Klein, M., Fischer, D., Schaar, A.C., Kuemmerle, L.B., Rybakov, S., Ibarra, I.L., Holmberg, O., Virshup, I., et al. (2022). Squidpy: a scalable framework for spatial omics analysis. Nat. Methods 19, 171–178. 10.1038/s41592-021-01358-2.

16. Pham, D., Tan, X., Xu, J., Grice, L.F., Lam, P.Y., Raghubar, A., Vukovic, J., Ruitenberg, M.J., and Nguyen, Q. (2020). stLearn: integrating spatial location, tissue morphology and gene expression to find cell types, cell-cell interactions and spatial trajectories within undissociated tissues. bioRxiv, 2020.05.31.125658. 10.1101/2020.05.31.125658.

17. Hao, Y., Hao, S., Andersen-Nissen, E., Mauck, W.M., Zheng, S., Butler, A., Lee, M.J., Wilk, A.J., Darby, C., Zager, M., et al. (2021). Integrated analysis of multimodal single-cell data. Cell 184, 3573-3587.e29. 10.1016/j.cell.2021.04.048.

18. Dries, R., Zhu, Q., Dong, R., Eng, C.-H.L., Li, H., Liu, K., Fu, Y., Zhao, T., Sarkar, A., Bao, F., et al. (2021). Giotto: a toolbox for integrative analysis and visualization of spatial expression data. Genome Biol. 22, 78. 10.1186/s13059-021-02286-2.

19. Wolf, F.A., Angerer, P., and Theis, F.J. (2018). SCANPY: large-scale single-cell gene expression data analysis. Genome Biol. 19, 15. 10.1186/s13059-017-1382-0.

20. Virshup, I., Bredikhin, D., Heumos, L., Palla, G., Sturm, G., Gayoso, A., Kats, I., Koutrouli, M., Community, S., Angerer, P., et al. (2023). The scverse project provides a computational ecosystem for single-cell omics data analysis. Nat. Biotechnol. 41, 604–606. 10.1038/s41587-023-01733-8.

21. Vizgen (2022). MERFISH FFPE Human Immuno-oncology Data Se.

22. Traag, V.A., Waltman, L., and Eck, N.J. van (2019). From Louvain to Leiden: guaranteeing well-connected communities. Sci. Rep. 9, 5233. 10.1038/s41598-019-41695-z.

23. He, S., Bhatt, R., Brown, C., Brown, E.A., Buhr, D.L., Chantranuvatana, K., Danaher, P., Dunaway, D., Garrison, R.G., Geiss, G., et al. (2022). High-plex Multiomic Analysis in FFPE at Subcellular Level by Spatial Molecular Imaging. bioRxiv, 2021.11.03.467020. 10.1101/2021.11.03.467020.

24. Efremova, M., Vento-Tormo, M., Teichmann, S.A., and Vento-Tormo, R. (2020). CellPhoneDB: inferring cell–cell communication from combined expression of multi-subunit ligand–receptor complexes. Nat. Protoc. 15, 1484–1506. 10.1038/s41596-020-0292-x.

25. Hunter, J.D. (2007). Matplotlib: A 2D Graphics Environment. Comput. Sci. Eng. 9, 90–95. 10.1109/mcse.2007.55.

26. Marcovecchio, P.M., Thomas, G., and Salek-Ardakani, S. (2021). CXCL9-expressing tumor-associated macrophages: new players in the fight against cancer. J. Immunother. Cancer 9, e002045. 10.1136/jitc-2020-002045.

27. Revel, M., Sautès-Fridman, C., Fridman, W.-H., and Roumenina, L.T. (2022). C1q+ macrophages: passengers or drivers of cancer progression. Trends Cancer 8, 517–526. 10.1016/j.trecan.2022.02.006.

28. Oft, M. (2014). IL-10: Master Switch from Tumor-Promoting Inflammation to Antitumor Immunity. Cancer Immunol. Res. 2, 194–199. 10.1158/2326-6066.cir-13-0214.

29. Bill, R., Wirapati, P., Messemaker, M., Roh, W., Zitti, B., Duval, F., Kiss, M., Park, J.C., Saal, T.M., Hoelzl, J., et al. (2023). CXCL9:SPP1 macrophage polarity identifies a network of cellular programs that control human cancers. Science 381, 515–524. 10.1126/science.ade2292.

30. Muntjewerff, E.M., Meesters, L.D., and Bogaart, G. van den (2020). Antigen Cross-Presentation by Macrophages. Front. Immunol. 11, 1276. 10.3389/fimmu.2020.01276.

31. Boutilier, A.J., and Elsawa, S.F. (2021). Macrophage Polarization States in the Tumor Microenvironment. Int. J. Mol. Sci. 22, 6995. 10.3390/ijms22136995.

32. Benner, B., Scarberry, L., Suarez-Kelly, L.P., Duggan, M.C., Campbell, A.R., Smith, E., Lapurga, G., Jiang, K., Butchar, J.P., Tridandapani, S., et al. (2019). Generation of monocyte-derived tumor-associated macrophages using tumor-conditioned media provides a novel method to study tumor-associated macrophages in vitro. J. Immunother. Cancer 7, 140. 10.1186/s40425-019-0622-0.

33. Ayers, M., Lunceford, J., Nebozhyn, M., Murphy, E., Loboda, A., Kaufman, D.R., Albright, A., Cheng, J.D., Kang, S.P., Shankaran, V., et al. (2017). IFN-γ–related mRNA profile predicts clinical response to PD-1 blockade. J. Clin. Investig. 127, 2930–2940. 10.1172/jci91190.

34. Gutiérrez-Melo, N., and Baumjohann, D. (2023). T follicular helper cells in cancer. Trends Cancer 9, 309–325. 10.1016/j.trecan.2022.12.007.

35. Tang, M., Kaymaz, Y., Logeman, B.L., Eichhorn, S., Liang, Z.S., Dulac, C., and Sackton, T.B. (2020). Evaluating single-cell cluster stability using the Jaccard similarity index. Bioinformatics 37, 2212–2214. 10.1093/bioinformatics/btaa956.

36. Grabski, I.N., Street, K., and Irizarry, R.A. (2023). Significance analysis for clustering with single-cell RNA-sequencing data. Nat. Methods 20, 1196–1202. 10.1038/s41592-023-01933-9.

37. Peyvandipour, A., Shafi, A., Saberian, N., and Draghici, S. (2020). Identification of cell types from single cell data using stable clustering. Sci. Rep. 10, 12349. 10.1038/s41598-020-66848-3.

38. Zappia, L., and Oshlack, A. (2018). Clustering trees: a visualization for evaluating clusterings at multiple resolutions. Giga Science 7, 1–9. 10.1093/gigascience/giy083.

39. Hart, T., and Zandbergen, P. (2014). Kernel density estimation and hotspot mapping. Polic.: Int. J. Polic. Strat. Manag. 37, 305–323. 10.1108/pijpsm-04-2013-0039.

40. Wand, M.P. (1994). Fast Computation of Multivariate Kernel Estimators. J. Comput. Graph. Stat. 3, 433–445. 10.1080/10618600.1994.10474656.

41. Tang, M. (2014). pyflow-RNAseq: a snakemake based RNAseq pipeline (Zenodo).

42. Köster, J., and Rahmann, S. (2012). Snakemake—a scalable bioinformatics workflow engine. Bioinformatics 28, 2520–2522. 10.1093/bioinformatics/bts480.

43. Dobin, A., Davis, C.A., Schlesinger, F., Drenkow, J., Zaleski, C., Jha, S., Batut, P., Chaisson, M., and Gingeras, T.R. (2013). STAR: ultrafast universal RNA-seq aligner. Bioinformatics 29, 15–21. 10.1093/bioinformatics/bts635.

44. Liao, Y., Smyth, G.K., and Shi, W. (2014). featureCounts: an efficient general purpose program for assigning sequence reads to genomic features. Bioinformatics 30, 923–930. 10.1093/bioinformatics/btt656.

45. Love, M.I., Huber, W., and Anders, S. (2014). Moderated estimation of fold change and dispersion for RNA-seq data with DESeq2. Genome Biol. 15, 550. 10.1186/s13059-014-0550-8.

