## Supplementary Figures for "Monkeybread: A Python toolkit for the analysis of cellular niches in single-cell resolution spatial transcriptomics data"

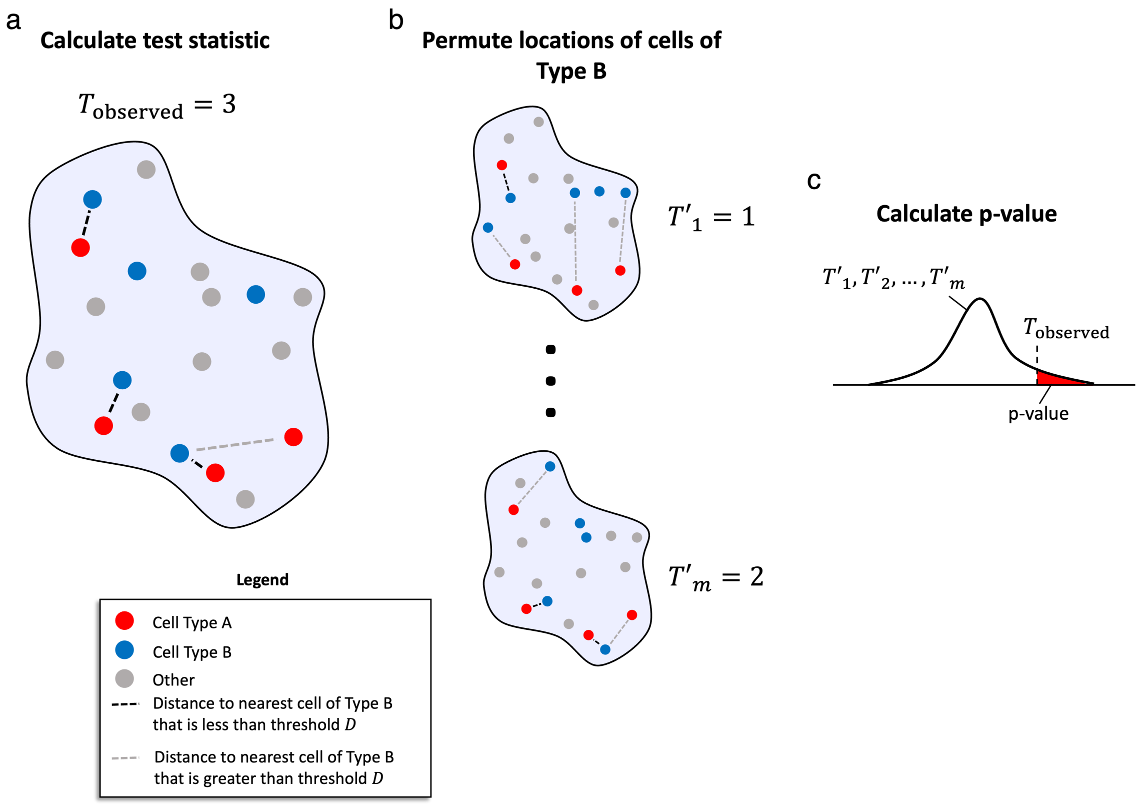


**Supplemental Figure 1. Overview of cell co-localization test. Related to Figure 2.** An overview of the cell co-localization test. **(a)** For every cell of type $A$, the distance to its nearest cell of type $B$ is computed. We then compute the test statistic by summing the number of these shortest distances than are less than a user-provided threshold. **(b)** Locations of non-type $A$ cells are permuted and the test statistic is re-calculated. **(c)** A permutation p-value is calculated.

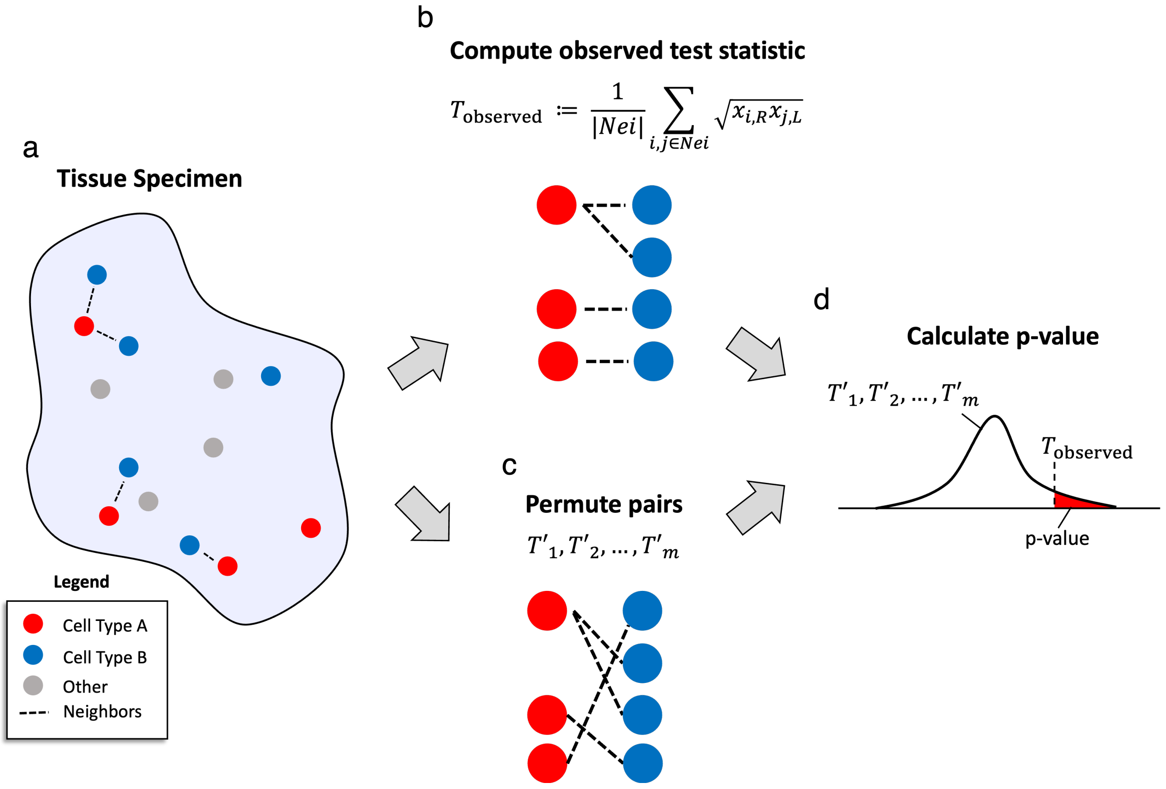


**Supplemental Figure 2. Overview of spatial ligand-receptor co-expression test. Related to Figure 2.** An overview of the ligand-receptor co-expression test described by He *et al*. (2021). **(a)** All pairs of neighboring cells between cell types $A$ and $B$ are determined based on their spatial proximities and **(b)** the ligand-receptor co-expression score is calculated. **(c)** The neighbor-pairings are permuted, and the ligand-receptor score is re-calculated for each permutation. **(d)** A permutation p-value is calculated.

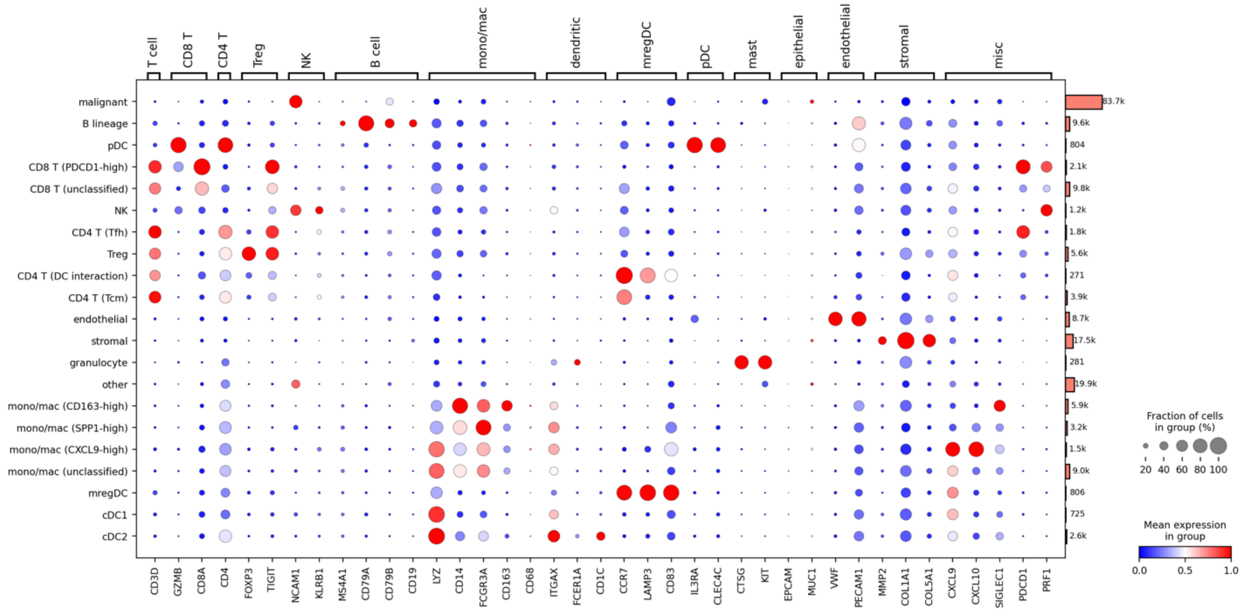


**Supplemental Figure 3. Marker gene expression within each cell subtype. Related to Figure 1.** A dot plot depicting expression of known marker genes within each annotated cell subtype in the human melanoma tumor dataset.

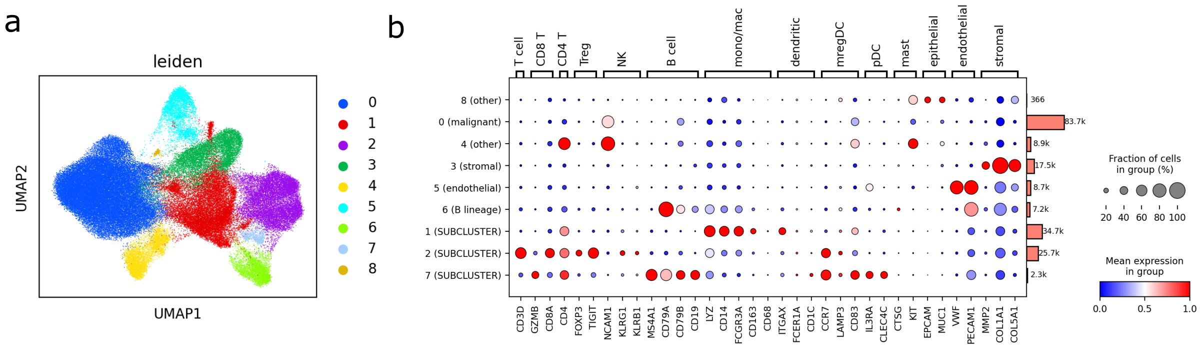


**Supplemental Figure 4. Clustering and annotation of all cells. Related to Figure 1. (a)** UMAP plot of human melanoma tumor cells colored by their cluster. For the sake of computational efficiency, a randomly subsampled half of the cells are shown. **(b)** A dot plot depicting expression of known marker genes within each cluster along with each cluster's annotation (in parentheses). Clusters labeled as "SUBCLUSTER" were subsequently subclustered and annotated.

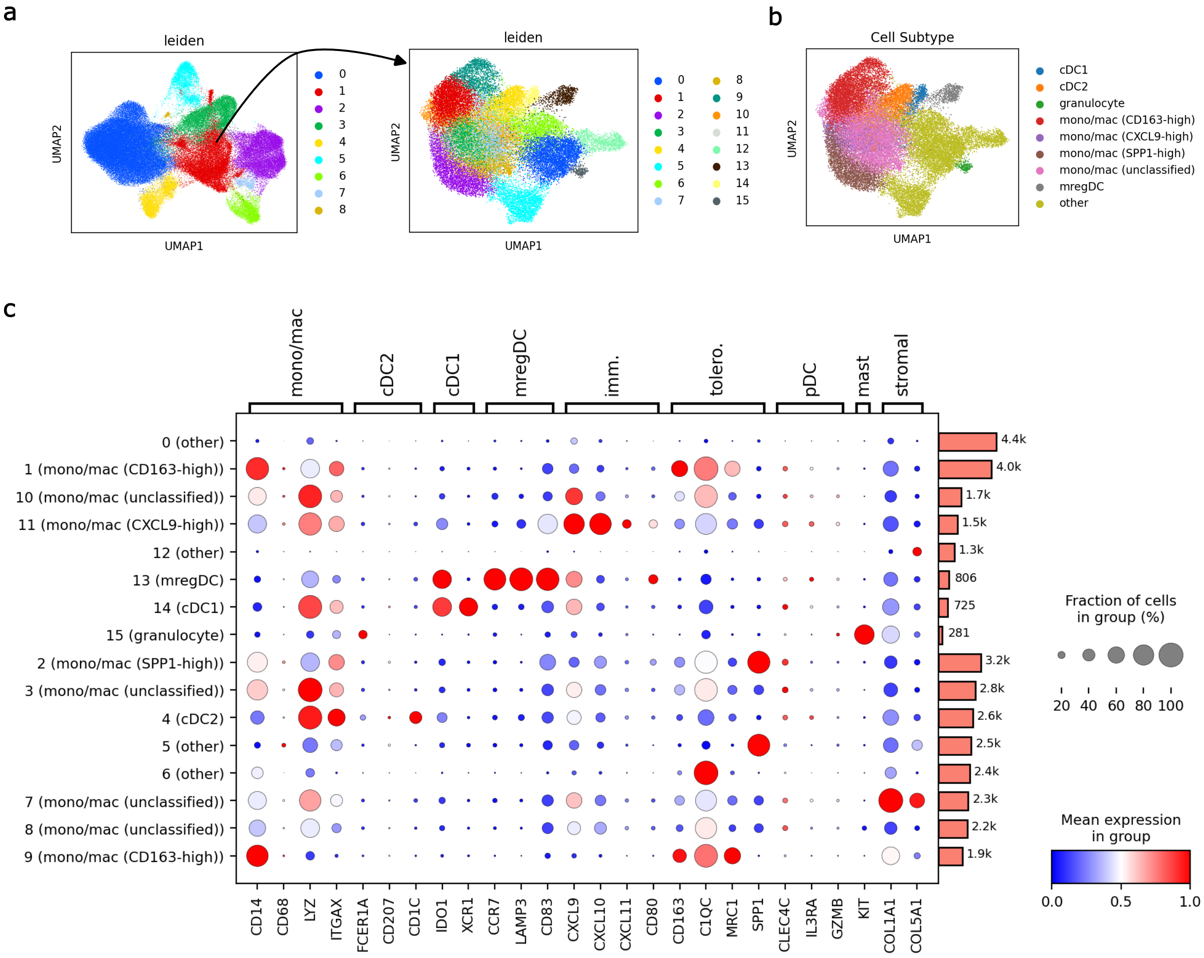


**Supplemental Figure 5. Subclusters of myeloid cells. Related to Figure 1. (a)** UMAP plot of cells from Cluster 1 of the human melanoma tumor cells colored by their subcluster and **(b)** annotated cell subtype. **(c)** A dot plot depicting expression of known marker genes within each subcluster along with each cluster's annotation (in parentheses).

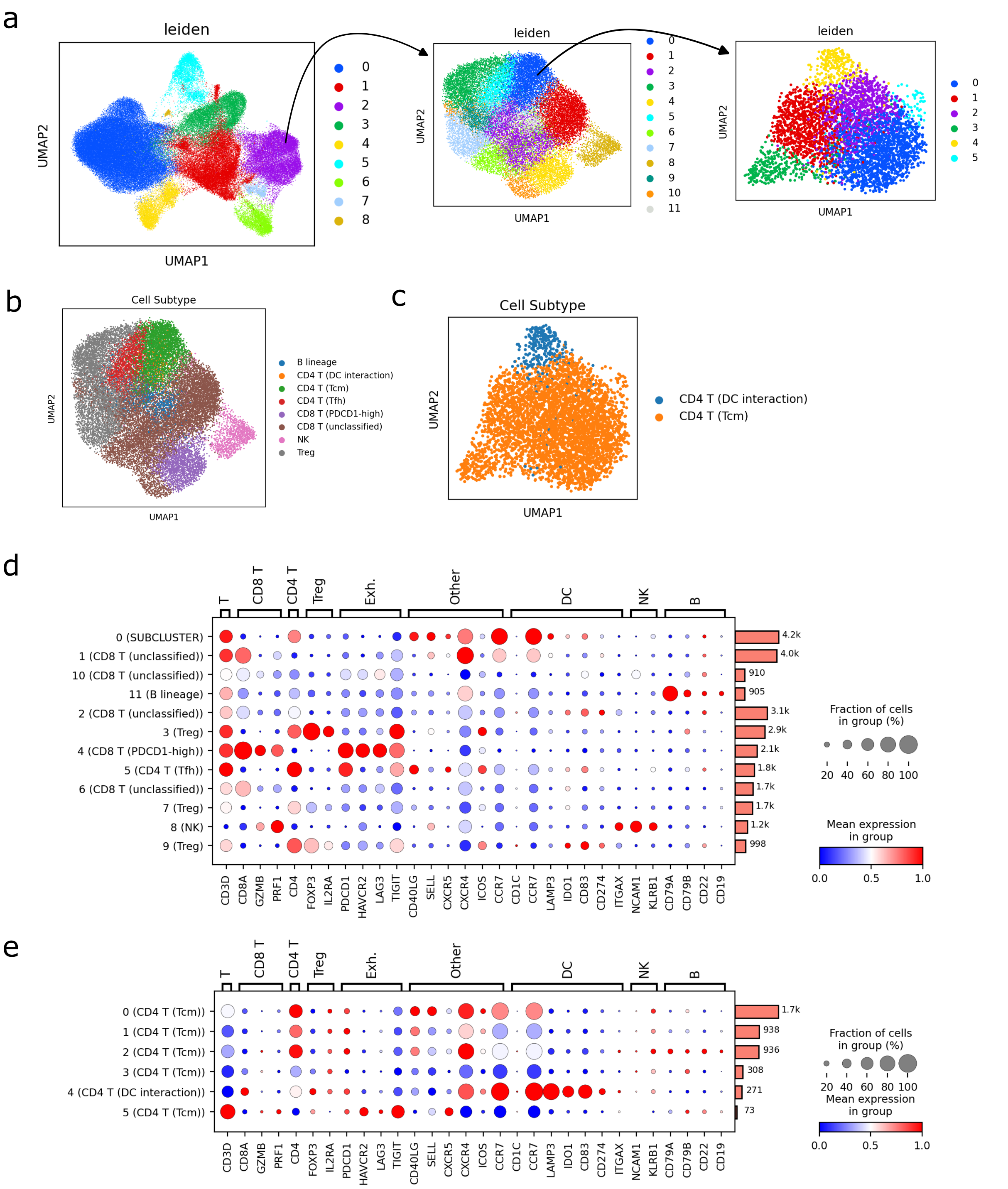


**Supplemental Figure 6. Subclusters of T cells. Related to Figure 1. (a)** UMAP plot of cells from Cluster 2 of the human melanoma tumor cells colored by their subcluster. Subcluster 0 was further subclustered. **(b)** UMAP plot of cell from Cluster 2, and **(c)** Subcluster 0 colored by their annotated subtype. **(d)** A dot plot depicting expression of known marker genes within each subcluster, and **(e)** each subcluster of Subcluster 0 along with the cluster's annotated cell type (in parentheses).

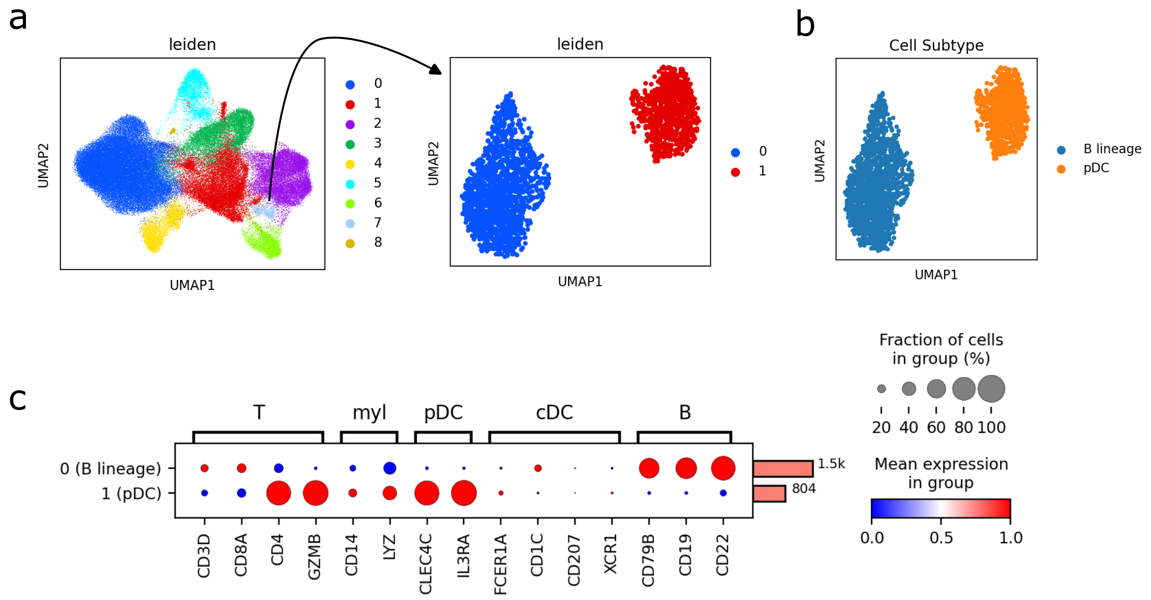


**Supplemental Figure 7. Subclusters of plasmacytoid dendritic cells and B cells. Related to Figure 1. (a)** UMAP plot of cells from Cluster 7 of the human melanoma tumor cells colored by their subcluster and **(b)** annotated cell subtype. **(c)** A dot plot depicting expression of known marker genes within each subcluster along with each cluster's annotation (in parentheses).
